# Fibroblast-derived HGF drives acinar lung cancer cell polarization through integrin-dependent RhoA-ROCK1 inhibition

**DOI:** 10.1101/185199

**Authors:** Anirban Datta, Emma Sandilands, Keith E. Mostov, David M. Bryant

## Abstract

The formation of lumens in epithelial tissues requires apical-basal polarization of cells, and the co-ordination of this individual polarity collectively around a contiguous lumen. Signals from the <u>*E*</u>xtra<u>*c*</u>ellular <u>*M*</u>atrix (ECM) instruct epithelia as to the orientation of where basal, and thus consequently apical, surfaces should be formed. We report that this pathway is normally absent in Calu-3 human lung adenocarcinoma cells in 3-Dimensional culture, but that paracrine signals from MRC5 lung fibroblasts can induce correct orientation of polarity and acinar morphogenesis. We identify HGF, acting through the c-Met receptor, as the key polarity-inducing morphogen, which acts to activate β1-integrin-dependent adhesion. HGF and ECM-derived integrin signals co-operate via a c-Src-dependent inhibition of the RhoA-ROCK1 signalling pathway via p190A RhoGAP. This occurred via controlling localization of these signalling pathways to the ECM-abutting surface of cells in 3-Dimensional culture. Thus, stromal derived signals can influence morphogenesis in epithelial cells by controlling activation and localization of cell polarity pathways.

## 1. Introduction

A simple epithelial monolayer consists of apical-basal polarized epithelial cells, which are the basic building block of acini in organs [1]. Epithelial cells possess functionally specialized basolateral and apical cell surfaces. The basolateral domain, although contiguous, is often described as two independent domains due to their distinct functional properties: the basal surface contacts the underlying <u>*e*</u>xtra<u>*c*</u>ellular <u>*m*</u>atrix (ECM) and the lateral surface allows adherence to neighbouring cells, while also providing a paracellular diffusion barrier [2]. In contrast, the apical surface provides the lining of the lumen of epithelial tissues, possessing the distinct characteristic of non-adherence to neighbours or the ECM. Such polarity on an individual level must be collectively coordinated between neighbouring cells to form the lumen and thus a biological tube.

While the mechanisms of how apical-basal polarity is formed have been extensively investigated [1, 2] how such polarity is oriented has until recently been neglected [3, 4]. Traditional culture methods for epithelial cells involves growth of cells to become apical-basal polarized monolayers on rigid substrata such as glass, plastic, or semi-permeable membrane filters. In these systems, the stiff culture vessel provides an immediate and strong cue as to where basal membranes should form [5]. In contrast, in 3-Dimensional (3D) culture where single epithelial cells are embedded in ECM gels, such as Matrigel or collagen, cells are surrounded by ECM. In such isotropic conditions, mechanisms must exist to orient apical-basal polarity formation such that the basal surface faces the ECM and the apical surface lines the cavity of ‘free space’.

We have previously described that temporospatial control of the RhoA signalling pathway is essential for the correct orientation of apical-basal polarity [6, 7]. In the MDCK cyst model, upon embedding isolated MDCK cells into Matrigel gels, single cells divide to form a cell doublet that has initially inverted polarity, i.e. apical surfaces erroneously facing the ECM. Detection of the ECM by β1-integrin-dependent adhesion results in phosphorylation of the RhoA-inhibitory GAP p190A RhoGAP [6]. P190A leads to inactivation of RhoA-Rho Kinase 1 (ROCK1) signalling, at the ECM abutting surface, allowing endocytosis of apical proteins from the ECM-abutting periphery and transcytosis to the centre of the cell doublet to form a lumen. This results in the correct orientation of apical-basal polarity between cells in 3D culture.

An open question from our and others’ previous studies is how generalizable signalling pathways identified to regulate polarity orientation in one cell type are to other epithelia? In addition, are these pathways defective in epithelial cancer cells? In the current work, we show that the previously identified pathway for correct polarity orientation is inactive in Calu-3 human lung adenocarcinoma cells grown in 3D at steady state. Fibroblast-derived Hepatocyte Growth Factor (HGF) can correct aberrant polarity orientation in 3D by controlling the localization of polarity pathways to the ECM-abutting surface. This suggests that future studies into modulation of this pathway may be important for understanding polarity changes in cancer.

## 2. Materials and Methods

### 2.1.2. Dimensional and 3-Dimensional culture

3D culture of Calu-3 was performed using adaptations of that described for MDCK [6]. Calu-3 (ATCC HTB-55) were grown in Eagle’s Minimum Essential Medium supplemented with 10% fetal bovine serum (FBS; Gibco). Cell line identity and lack of mycoplasma was confirmed (CRUK Beatson Institute). For plating in 3D, single cell suspensions were made (2 × 10^4^ cells ml^−1^) in growth medium supplemented with 2% Matrigel (BD Biosciences). Cell:medium:Matrigel mixture was plated onto Matrigel-pre-coated (15μl of net Matrigel) 8-well coverglass chambers (Nunc, LabTek-II). At appropriate times paraformaldehyde (PFA) was used to fix cells. For induction of acinar morphogenesis, cells were treated at day 4 with modifiers and fixed at day 9. Modifiers included MRC5 lung-derived fibroblast-conditioned medium (FCM), HGF blocking antibody (0.1 μg/mL; R&D Systems, 24612), SU11274 (10 μM; Tocris), HGF (10 ng/ml, gift from Genentech), AIIB2 (1:200; a gift of C. Damsky, UCSF), TS2/16 (1:100; a gift of C. Damsky, UCSF), PP2 (5 μM, Sigma-Aldrich, P0042), and Y-27632 (10 μM, Calbiochem, 688002). MRC5 (ATCC) were cultured in Eagle’s Minimum Essential Medium supplemented with 10% fetal bovine serum (FBS; Gibco). For production of condition medium MRC5 were grown as previously described [8] to confluence in Eagle’s Minimum Essential Medium supplemented with 10% fetal bovine serum, 100 U/ml penicillin, 100 mg/ml streptomycin, in tissue culture flasks (T75; Corning). Fresh medium was replaced (30 ml) before culture for 3 days. Fibroblast conditioned medium was centrifuged at 3,500 & g to remove cell debris, and frozen at −20°C.

### 2.2. Antibodies and immunolabelling

Calu-3 in 3D culture were stained essentially as described for MDCK cysts [9]. Primary antibodies utilized were: Ki-67 (Epitomics, 2642-1), Cleaved Caspase 3 (Cell Signaling Technology, 9661L), JAM-A (BD Biosciences, 612120), Muc1 (Epitomics, 2900-1), GM130 (BD Biosciences, C65120), β-catenin (Santa Cruz, sc-7199), NHERF1 (Abcam, ab3452), Ezrin (BD Biosciences, 610603), Cleaved Caspase 8 (Cell Signaling Technology, 9496), β1-integrin (BD Biosciences, 610467), pY416-c-Src (Cell Signaling Technology, 6943P), pY1105-p190 (Abcam, ab55339). Alexa fluorophore-conjugated secondary antibodies (1:250) or Phalloidin (1:200) (both Invitrogen) and Hoechst to label nuclei (10 μg/ml), were utilized. Imaging was performed on a Zeiss 510 Confocal Microscope, using a 63x oil immersion lens. Image processing was performed using ImageJ. Images shown are representative of 3 separate experiments.

### 2.3. Immunoblotting

Protein blotting was performed as described [6]. To solubilize cells, cultures were washed twice in ice-cold PBS, before addition of ice-cold extraction buffer [50 mM Tris-HCl pH 7.4, 150 mM NaCl, 0.5 mM MgCl_2_, 0.2 mM EGTA, and 1% Triton X-100 plus 50 mM NaF, 1 mM Na_3_VO_4_ and complete protease inhibitor cocktail tablet (Roche, Mannheim, Germany)]. Lysates were extracted at 4°C for 25 min. For 3D cultures a 27½-gauge needle was used for assisting extraction from ECM. To remove debris, centrifugation at 14,000 × *g* at 4°C for 10 min was performed. Samples were separated using SDS-PAGE, and were transferred to PVDF membranes. Western analysis was performed using either chemiluminescence (SuperSignal Chemiluminescence Kit; Pierce, Rockford, IL) or infrared fluorescent secondary antibodies and quantitative detection (Odyssey CLx, Li-COR Biosciences). A BCA Protein Assay Reagent kit (Pierce) was used to determine protein concentration. Transfer and protein loading were monitored by staining gels with 0.1% Coomassie Brilliant Blue. Antibodies for WB were: GAPDH (Millipore, MAB374), pY416-c-Src (Cell Signaling Technology, 6943P), total c-Src (Cell Signaling Technology, 2102P), pY1105-p190 (Abcam, ab55339), P190A (BD Biosciences, 610149), RhoA (Santa Cruz, sc-418), Phospho-Myosin Light Chain 2 (Ser19; Cell Signaling Technology, 3671L), ROCK1 (Santa Cruz, sc-17794), ROCK2 (Santa Cruz Biotechnology, sc-100425).

### 2.4. RNAi, virus production and transduction

Stable depletion of proteins was performed using pLKO.1-puro lentiviral shRNA vectors, obtained from the TRC collection. Sequences and clone identities are as follows: shScramble (5’-CCTAAGGTTAAGTCGCCCTCG-3’) [10], shc-Src_1 (TRCN0000038149, 5’-GCTCGGCTCATTGAAGACAAT-3’), shc-Src_2 (TRCN0000038150, 5’-GACAGACCTGTCCTTCAAGAA-3’), p190ARhoGAP_1 (TRCN0000022188, 5’-CGGTTGGTTCATGGGTACATT-3’), p190ARhoGAP_2 (TRCN0000022184, 5’-GCCCTTATTCTGAAACACATT-3’), shRhoA_1 (TRCN0000047710, 5’-GTACATGGAGTGTTCAGCAAA-3’), shRhoA_2 (TRCN0000047711, 5’-TGGAAAGACATGCTTGCTCAT-3’), shROCK1_1 (TRCN0000121094, 5’-CGGTTAGAACAAGAGGTAAAT-3’), shROCK1_2 (TRCN0000121095, 5’-GCATTCCAAGATGATCGTTAT-3’), shROCK2_1 (TRCN0000000978, 5’-GCACAGTTTGAGAAGCAGCTA-3’), shROCK2_2 (TRCN0000000981, 5’- CCTCAAACAGTCACAGCAGAA-3’).

To produce lentiviral particles, pLKO plasmids were co-transfected with Virapower (Invitrogen) lentiviral packaging mix into packaging cells according to manufacturer instructions (297-FT, Invitrogen). At time of collection, viral supernatants were centrifuged twice at 3500 × g to remove cell debris. Cells were transduced with lentiviruses the day after plating, and were infected for 60 hours, before selection with puromycin (5 μg/ml, Life Technologies, A11138-03). Efficient depletion was verified using Western blotting.

### 2.5 Adhesion Assay

To determine adhesion capability to laminin, 96 well plates were coated with 10ug/ml laminin-1 (BD, 354239) in PBS+ at 37°C for 30 min. Wells were washed with PBS to remove non-bound laminin-1 before blocking of non-coated regions with 1% BSA solution (SigmaAldrich) in PBS+ for 30 min. Isolated Calu-3 cells were seeded at 10,000 cells per well of a 96-well plate (100,000 cells/ml in 100ml per well) in adhesion buffer (HBSS, 10mM HEPES, 2mM Glucose) and incubated for 1 hour. Wells were gently washed with PBS to remove unbound cells. Adhesion to the well was determined by staining with Calcein AM (Life Technologies) for 5 min, and reading for fluorescence in a microplate reader.

### 2.6 FACS Analysis

For quantitation of surface β1-integrin levels, cells were washed twice with PBS, trypsinised, centrifuged at 1,500 RPM then re-suspended in 500 μl of FACS buffer (PBS+0.1% heat inactivated BSA) and kept on ice. For primary antibody labelling with β1-integrin (BD Biosciences, 610467), cells were centrifuged and re-suspended in 100 μl of FACS buffer containing a 1:200 dilution of primary antibody and incubated on ice for 30 min. Cells were washed twice by sequential addition of 1 ml FACS buffer followed by centrifugation as above. Alexa 488 secondary antibody (Thermo Scientific) was added according to the antibody staining as above. Two rounds of washing were performed before surface integrin levels were measured using a FACS Aria 2 (BD).

### 2.7 Statistical tests

Quantitation of 3D acinus formation was performed as adapted from MDCK cultures as previously described [9]. The percentage of acini displaying a single central lumen was determined, and compared to control cell aggregates. Values are mean ± s.d. from three experiments, with *n* ≥100 cysts per experiment. Significance was calculated using a paired, two-tailed Student’s *t*-test.

## 3. Results

### 3.1 Fibroblast-Conditioned Medium induces acinar polarization of Calu-3 in 3-Dimesional culture

We examined the morphogenesis of a lung adenocarcinoma cell line Calu-3 in 3-Dimensional (3D) culture, which is thought to be derived from a tumour of the bronchial submucosal glands [11]. 3D culture of Calu-3 for up to 9 days in standard growth medium (Methods and Materials) failed to induce acinar morphogenesis, forming solid multicellular aggregates (Fig. 1A, B). We next examined, the influence of soluble factors secreted by an established lung stromal fibroblast cell line (MRC5) on Calu-3 3D morphogenesis. We cultured Calu-3 cells for 9 days, modifying the medium by the addition of various factors from days 4-9 (Fig. 1A). Strikingly, morphogenetic rearrangements of the Calu-3 cells, resulting in lumen-containing acini, could be optimally achieved by changing the culture medium to MRC5 lung fibroblast-conditioned medium (FCM) from day 4-9 (Fig. 1A-B). Addition of FCM at earlier time points resulted in single cell invasion (data not shown), suggesting that aggregates needed to reach a certain size before rearrangement of polarity could occur. We thus used this 4+5 day schedule for subsequent experiments.

**Figure 1:**
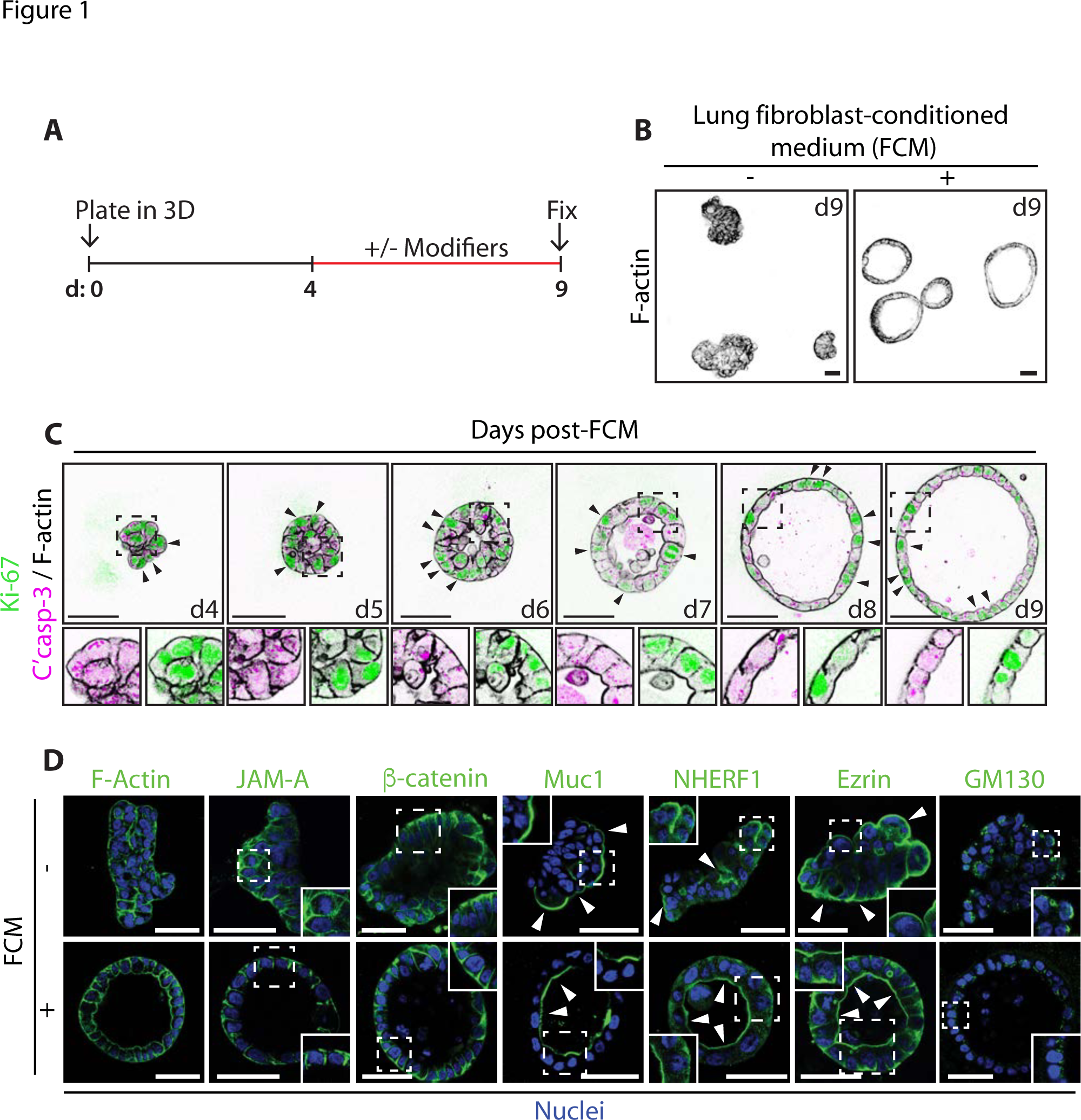
Fibroblast Conditioned Medium (FCM) induces the formation of polarized Calu-3 acini. (A) Schematic showing the experimental procedure for developing Calu-3 acini in 3D with key modifiers added or removed between days 4-9. (B) Calu-3 cells were cultured on a layer of Matrigel in standard growth medium containing 2% Matrigel. After 4 days the media on the resultant aggregates were either replenished or replaced with FCM. Acini were fixed on day 9, stained with F-actin and imaged using a confocal microscope. (C) Acini were fixed and stained at different time points post addition of FCM at day 4, with anti-Ki-67 (green), or anti-Cleaved Caspase-3 (magenta) antibodies and with Phalloidin (F-actin, black). Arrows indicate the localization of Ki-67. Magnified images are also shown (lower panels). (D) Untreated aggregates or FCM-treated acini were fixed and stained with Hoechst (Nuclei, blue) and either Phalloidin (F-actin), or anti-JAM-A, anti-β-catenin, anti-Muc1 anti-NHERF-1, anti-Ezrin or anti-GM130 antibodies (all green). Arrows indicate the localization of apical proteins. Insets show magnified images. All scale bars, 50 μm.

The first notable event in FCM-induced morphogenesis was the transition of cell aggregates from a non-spherical cluster into a sphere by day 5 (d5) (Fig. 1C). Subsequently, cells not rearranged into the acinus wall were cleared, concomitant with the appearance of a central lumen, and resulting in a cleared and expanded lumen by d9. Examination of proliferation (Ki-67) and apoptosis (cleaved caspase-3) markers in response to FCM revealed a redistribution of proliferation markers from all cells (d4) to solely the acinus wall (d9), while apoptosis could be observed in the lumen (d6-8; Fig 1C). FCM-treated acini possess luminal surface localization of the apical proteins Muc1, NHERF1 and Ezrin, apically oriented Golgi, and basolateral surface β-catenin and JAM-A (Fig. 1D). Without FCM, basolateral protein and Golgi orientation appeared randomized, while apical proteins were oriented towards the ECM (Fig. 1D). This suggests that FCM caused a polarity reversion accompanied by formation of polarized acini.

### 3.2 HGF is the morphogen in FCM inducing acinar morphogenesis

We aimed to determine the morphogenetic factor in FCM that induce Calu-3 acinar morphogenesis. Hepatocyte Growth Factor (HGF) is a key morphogen that MRC5 secrete to induce tubulogenesis of MDCK cysts [8, 12]. Surprisingly, addition of HGF alone was sufficient to induce acinus formation by Calu-3 (Fig. 2A, B), completely mirroring the effects of FCM. Moreover, signalling by HGF and its receptor, c-Met, were both necessary for acinus formation by Calu-3, as addition of c-Met inhibitor or HGF blocking antibodies abolished the effects of FCM (Fig. 2A, C). HGF-induced morphogenesis mirrored that observed for FCM, converting non-apoptotic cell aggregates into polarized acini with lumens that display apoptotic remnants, as indicated by the presence of cleaved caspase 3/8 in the lumens of acini (Fig. 2B). These data suggest that HGF/c-Met signalling is a key pathway activated in Calu-3 by FCM to regulate Calu-3 acinus formation.

**Figure 2:**
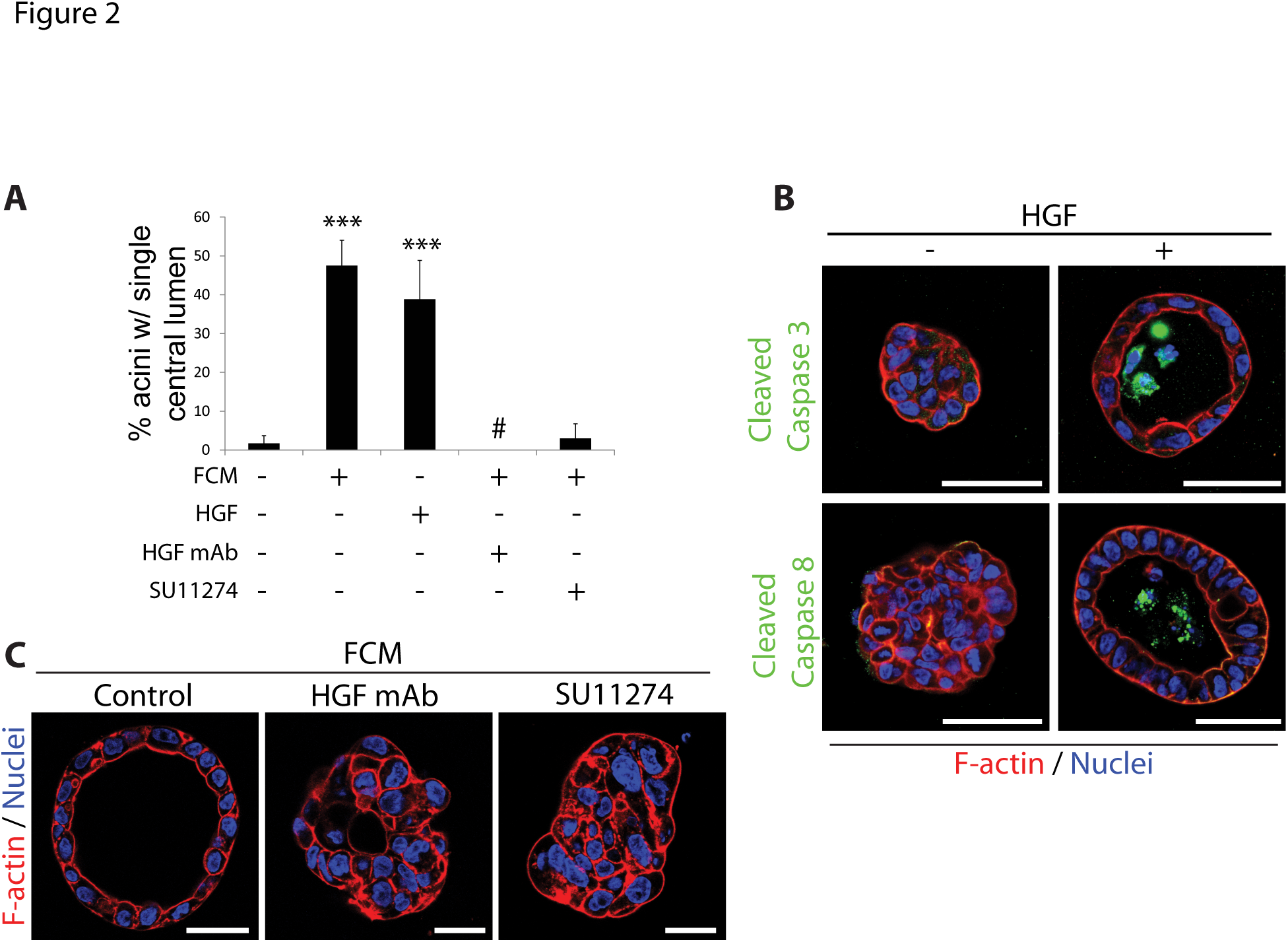
HGF stimulate Calu-3 acinus formation. (A) Calu-3 aggregates were treated on day 4 with FCM, HGF (10 ng/ml), FCM + HGF blocking monoclonal antibody (HGF mAb; 0.1 μg/mL) or FCM + c-Met Inhibitor (SU11274; 10 μM) and fixed on day 9. Graph shows the number of acini with a single central lumen. Results are presented as mean percentage +/- s.d. and significance is ***p≤0.0001 (n=3 experiments). (B) Aggregates/acini formed in the absence or presence of HGF were fixed and stained with either anti-Cleaved Caspase 3 (green in upper panels) or anti-Cleaved Caspase 8 antibodies (green in lower panels) and F-actin (red) and Hoechst (blue). (C) Calu-3 aggregates were treated on day 4 with FCM, FCM + HGF blocking monoclonal antibody (HGF mAb; 0.1 μg/mL) or FCM + c-Met Inhibitor (SU11274; 10 μM) and fixed on day 9. Staining was performed for F-actin (red) and Nuclei (Hoechst, blue). All scale bars, 50 μm.

### 3.3 Acinar morphogenesis involves β1-integrin-dependent adhesion and c-Src kinase

MDCK cells grown in 3D can form either apical-basal polarized cysts surrounding a central lumen, or can be induced to form front-rear polarized structures with inverted polarity by perturbing signalling from the ECM [9]. We examined whether HGF modulates ECM signalling in Calu-3 cells. In 2D culture conditions on plastic, Calu-3 grow as loosely adherent aggregates (Fig. 3A). Addition of HGF in 2D resulted in morphogenetic rearrangement whereby cells now flattened and spread on the plastic substratum. We measured adhesion of freshly plated cells to the major ECM component of Matrigel, laminin-1, and found that in contrast to the normally low adhesive index of Calu-3, HGF induced strong adhesion of cells (Fig. 3B). This could be completely blocked by inhibition of β1-integrins, as could steady state adhesion levels. Strikingly, activating antibodies to β1-integrins alone were sufficient to induce adhesion of Calu-3, to levels above that of HGF alone, and could further enhance the effect of HGF (Fig. 3B). These data suggest that HGF may act to induce morphogenesis by activating β1-integrin signalling.

**Figure 3:**
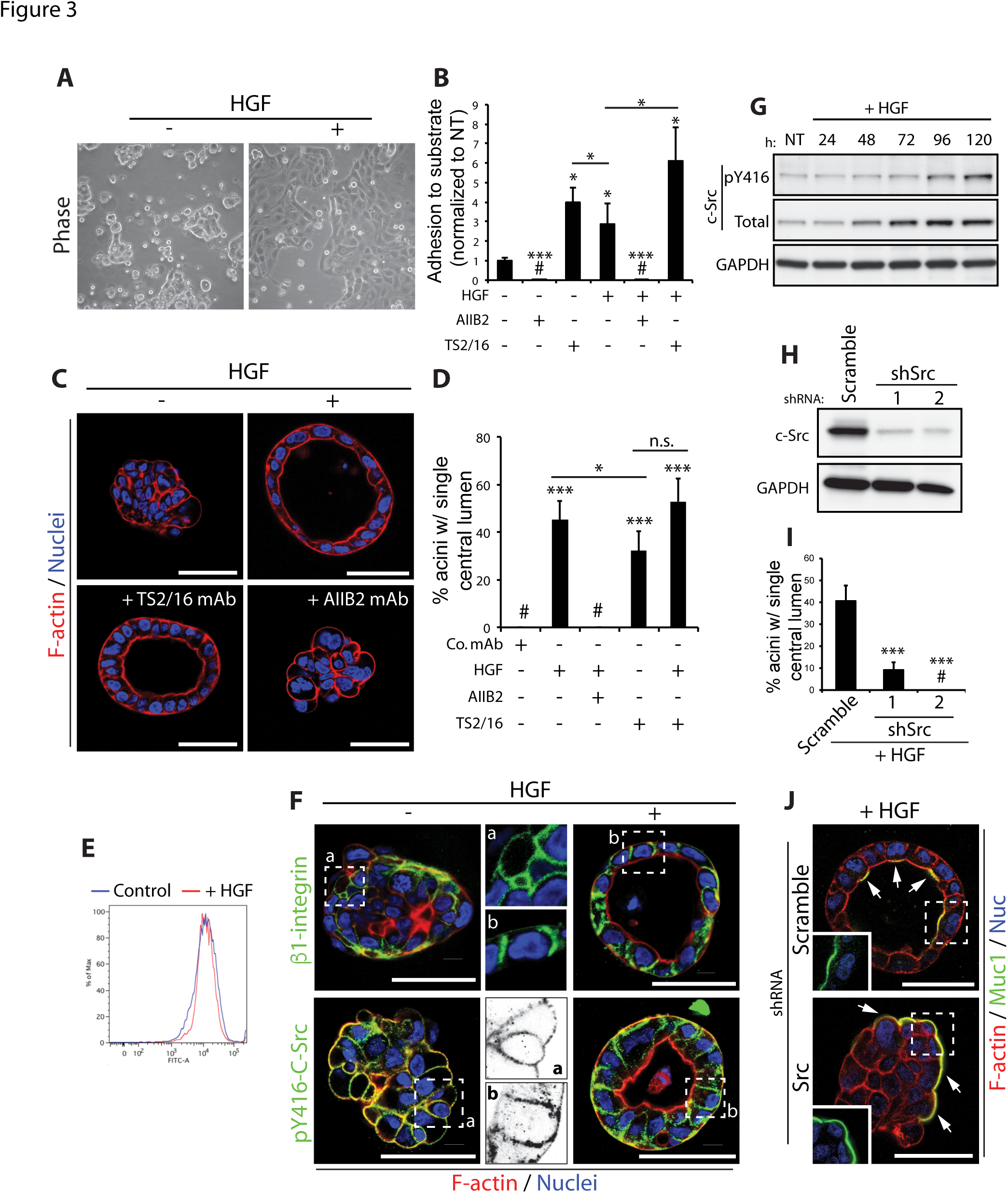
c-Src is a major target of HGF signalling in acinar morphogenesis. (A) Phase images of Calu-3 cells grown on plastic in presence or absence of HGF. (B) Graph shows the relative adhesion of Calu-3 cells adhering to laminin-1 after treatment with HGF and either AIIB2, a β1 integrin blocking antibody (1:200), or TS2/16, a β1 integrin activating antibody (1:100), all normalized to non-treated control cells (NT). (C) Control aggregates or aggregates treated with HGF, with TS2/16 or with HGF and AIIB2, were fixed and stained with F-actin (red) and Hoechst (blue). (D) Graph shows the percentage of acini with a single central lumen upon treatment with either control antibody (Co. mAb), HGF, AIIB2 or TS2/16. (E) Surface expression of β1-integrin in control and HGF treated Calu-3 cells in 2Dwas examined by FACS analysis. (F) Untreated aggregates or HGF-treated acini were fixed and stained with anti-β1integrin (green in upper panels) or anti-pY416-c-Src (green in lower panels)antibodies and F-actin (red) and Hoechst (blue). A magnified image of anti-β1integrin (upper middle panels) and anti-pY416-c-Src (lower middle panels) in untreated and HGF treated samples is also shown (a and b respectively). (G) Aggregates were stimulated with HGF and harvested at various time-points.Immunoblotting was performed using anti-pY416-c-Src, anti-c-Src and anti-GAPDH antibodies. (H) c-Src protein expression was reduced in Calu-3 cells using either of 2 different shRNAs and knockdown confirmed by immunoblotting with anti-c-Src and anti-GAPDH antibodies. (I) Graph shows the percentage of acini with a single central lumen after either control of c-Src knockdown and upon HGF stimulation. (J) Aggregates expressing Scramble or c-Src shRNA were stimulated with HGF, fixed and stained with anti-Muc1 antibody (green), F-actin (red) and Hoechst (blue). Arrows indicate localization of Muc1 at membranes. Insets show magnified images of Muc1 localization (green). Graphs (except 3B) are presented as mean percentage +/- s.d. and significance is *p≤0.05 and ***p≤0.0001 (n= 3). All scale bars, 50 μm.

β1-integrin similarly controlled 3D morphogenesis. Inhibiting β1-integrins in 3D completely blocked the effect of HGF on acinar morphogenesis. Strikingly, activating β1-integrin alone was sufficient to induce acinar morphogenesis (Fig. 3C, D). In 3D, combined HGF treatment and β1-integrin experimental activation failed to cause further acinar morphogenesis, suggesting that each treatment may maximally activate β1-integrins in this system. These data suggest that β1-integrins signalling pathway may be inactive in Calu-3 at steady state, and activation by HGF or antibodies is sufficient to restore 2D adhesion and 3D acinus formation.

We examined the mechanism of HGF-induced β1-integrin function. Using flow cytometry from cells from 2D, HGF simulation did not alter the amount of β1-integrins on the surface (Fig. 3E), suggesting that presence vs. absence of cortical integrin does not account for the effects of HGF. In 3D, β1-integrin localized to the cell cortex in an apparently unpolarised fashion. In contrast, HGF induced a rearrangement of polarity such that β1-integrin now localized only to basolateral membranes (Fig. 3F). These data suggest that HGF induces cell polarity by triggering the cortical asymmetry of β1-integrins solely to the basolateral membrane-ECM interface to prompt acinus formation.

To investigate this further we examined a common signalling effector of β1-integrins, the kinase c-Src [13]. In 2D, HGF stimulation induced activation of c-Src, which could be blocked by either a c-Src inhibitor or by blocking β1-integrins (Fig. S1A). In 3D, HGF induced a somewhat delayed activation of c-Src, concomitant with an increase in total c-Src levels (Fig. 3G). We noted polarization of activated c-Src (pY416-c-Src; p-c-Src) similar to β1-integrin in response to HGF, moving from general cortical localization to exclusively basolateral membranes (Fig. 3F). c-Src was required for HGF induced morphogenesis, as chemical inhibition or the depletion of c-Src with either of two shRNAs resulted in a robust inhibition of acinus formation in response to HGF (Fig. 3H-J, S1B). c-Src is thus a major target of HGF signalling in acinar morphogenesis.

### 3.4 Inhibition of the RhoA-ROCK1 signalling pathway is required for acinar polarization

In MDCK cysts, β1-integrin signalling inhibits apical domain formation at the ECM-abutting cyst periphery by promoting peripheral phosphorylation of the RhoA-specific inactivator p190A RhoGAP (hereforth referred to as p190A) [9]. In Calu-3 in 3D, HGF/c-Src signalling regulated p190A re-localization rather than controlling p190A phosphorylation (Fig. 4A, B). In control aggregates, p190A localized to cell-cell contacts, but was absent from ECM-abutting membranes. HGF induced p190A re-localization exclusively to the basolateral domain (Fig. 4A), mirroring that observed for β1-integrin and pY416-c-Src (Fig. 3). P190A was required for HGF-induced acinus formation, as p190A depletion resulted in highly disorganized aggregates displaying inverted polarity despite HGF stimulation (Fig. 4C-E). This could be rescued by inhibition of Rho Kinase (ROCK1/2) (Fig. 4E), confirming that p190A is acting as an inhibitor of the Rho-ROCK pathway to control polarity orientation.

**Figure 4:**
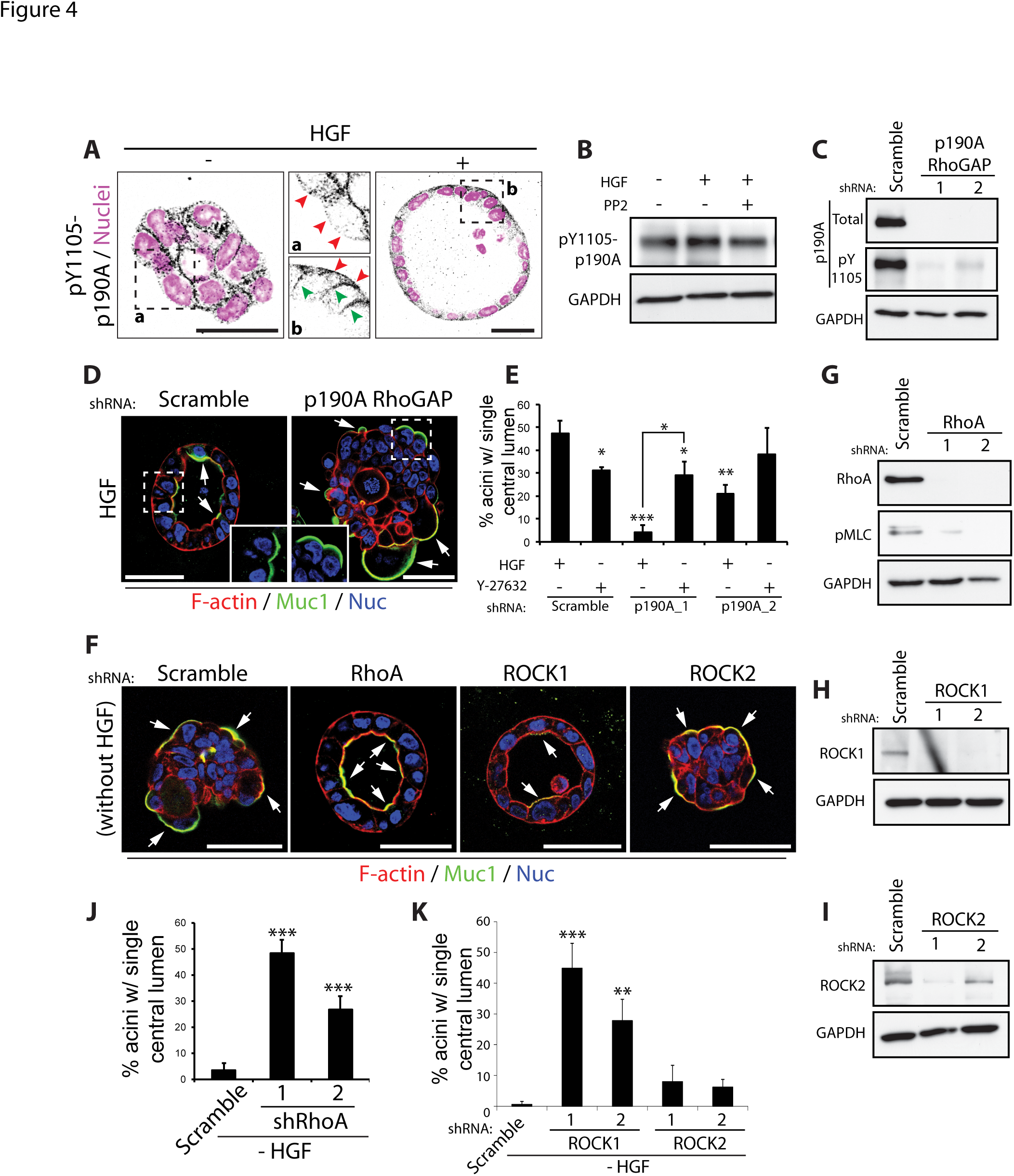
A RhoA-ROCK1 inhibitory module downstream of c-Src and β1 integrin controls acinus formation. (A) Aggregates were treated with HGF, fixed and stained with pY1105-p190ARhoGAP antibody (black) and Hoechst (magenta). Magnified images of pY1105-p190A RhoGAP localization in untreated and HGF stimulated samples (middle panels, a and b respectively) are shown. Red arrows indicate presence or absence of pY1105-p190A RhoGAP at the ECM-abutting surface, while green arrows show its localization to lateral membranes. (B) Aggregates treated with HGF and the c-Src inhibitor PP2 (5 μM), were harvested and immunoblotted with anti-pY1105-p190A RhoGAP and anti-GAPDH antibodies. (C) P190A RhoGAP expression was reduced in Calu-3 cells using either of 2 different shRNAs and knockdown confirmed by immunoblotting with anti-p190A RhoGAP, anti-pY1105-p190A RhoGAP and anti-GAPDH antibodies. (D) Calu-3 aggregates expressing Scramble or p190A RhoGAP shRNA were stimulated with HGF then fixed and stained with anti-Muc1 antibody (green), F-actin (red) and Hoechst (blue). Arrows indicate localization of Muc1 at membranes. Insets show magnified images of Muc1 localization (green). (E) Graph shows the percentage of acini with a single central lumen after treatment with HGF and Rho Kinase (ROCK1/2) inhibitor, Y-27632 (10 μM). (F) RhoA, ROCK1 and ROCK2 were depleted in Calu3 cells using specific shRNAs and samples stained with anti-Muc1 antibody (green), F-actin (red) and Hoechst(blue). Arrows indicate localization of Muc1 at membranes. (G-I) Knockdown with either of 2 different shRNAs was confirmed by immunoblotting with (G) anti-RhoA, (H) anti-ROCK1 and (I) anti-ROCK2 antibodies. GAPDH antibody was used as a control. (J-K) Graphs show the percentage of acini with a single central lumen after knockdown of (J) RhoA or (K) ROCK1 and ROCK2. All graphs are presented as mean percentage +/- s.d. and significance is *p≤0.05, **p≤0.001 and ***p≤0.0001 (n= 3). All scale bars, 50 μm.

Our data reveal that suppression of the ROCK signalling pathway may be sufficient to promote acinus formation in 3D Calu-3. Depletion of RhoA and ROCK1, but not ROCK2, was sufficient to induce acinus formation, abolishing the requirement for HGF (Fig. 4E-K). ROCK inhibition alone was slightly more potent than HGF at inducing acinar polarisation, and combined HGF treatment with ROCK inhibition had an additive effect on acinus formation (Fig. S1C). These data suggest that HGF activates a c-Src-and β1-integrin-dependent pathway that converges upon a RhoA-ROCK1 inhibitory module to control acinus formation.

Together, our data reveal an HGF:c-Met-dependent activation of integrin-based adhesion to regulate polarity through c-Src:p190A-dependent inhibition of RhoA-ROCK1 signalling. As a similar pathway exists in MDCK cysts [6], and the correct orientation of polarity can be ‘rescued’ in organoid cultures from patients displaying inverted polarity [14], our data suggest a conserved role for p190A, RhoA, and ROCK1 in controlling epithelial polarity orientation.

## 4. Discussion

In this study, we set out to examine the mechanisms of apical-basal polarity orientation pathways in 3D culture. We identified a highly similar pathway of polarity orientation in Calu-3 acini as MDCK cysts culture, indicating a conservation of polarity mechanisms [6]. In both systems, β1-integrin is required to localize a kinase to basolateral membranes to phosphorylate p190A RhoGAP at a site, which brings it into a complex with β1-integrins. c-Src and FAK are central to this in both MDCK and Calu-3 acini (unpublished observations). This sets up a zone of inhibition against the RhoA GTPase at the ECM-abutting membrane of cells, resulting in the inhibition of RhoA and its downstream kinase ROCK1. This collectively allows the reorientation of apical-basal polarity to form a lumen.

Most studies examining epithelial polarity almost exclusively utilize isolated epithelial cells [15]. Even in such systems the regulation of polarity orientation is only recently becoming clear. In Calu-3 cells in 3D, though the polarity-regulating pathway exists and localises to the cell cortex, it does not display the required asymmetry – solely to the basolateral domain – to facilitate cell polarisation. This requires activation by HGF/c-Met signalling, which can occur upon paracrine signalling by fibroblast-derived HGF. The cross-talk between stromal and epithelial compartments in a tissue are essential for morphogenesis of both compartments *in vivo* during development and in homeostasis. For instance, during kidney development the branching of the ureteric bud is induced by the neighbouring stromal compartment, the metanephric mesenchyme, which in turn feeds back on the mesenchyme to promote condensation of the stroma to become lumen-containing nephron precursors [16]. In addition, in cancer progression, tumours can influence the neighbouring stroma, activating it to become cancer-associated fibroblasts, which again feedback to promote the progression of tumourigenesis [17]. Such bidirectional cross talk between the stroma and the epithelium suggests that overall tissue architecture is an emergent property of many cell types, each influencing the other to regulate collective organisation. It will thus be important in future studies to examine co-culture of epithelia with other cell types to determine how these influence cell polarity.

The RhoA-ROCK1 signalling pathway is a major regulator of polarity orientation [3, 4, 6, 7], and our current and previous results indicate that it must be under strict spatiotemporal control to allow for the correct orientation of apical-basal polarization. Inhibition of RhoA-ROCK1 signalling appears to be a common requirement for robust polarization in 3D contexts, for a growing list of cell types, including MDCK cysts [6, 7, 18], Calu-3 as indicated here, and explants of primary human intestinal cells [14]. Moreover, ROCK inhibitor is a common component of most medium required to establish *ex vivo* organoid growth and polarisation around a central lumen [15, 19]. Whether p190A RhoGAP is the main regulator of this, or whether there are many layers of redundancy to ensure inhibition of RhoA-ROCK1 signalling is unclear. Inhibition of RhoA-ROCK1 does appear to be a generalized requirement for efficient lumen formation.

Given the largely overlapping targets of ROCK1 and ROCK2 [20], it is surprising that only ROCK1 inhibition is responsible for polarity reorientation. Our understanding of how ROCK1 antagonises luminal polarisation is incomplete. In the MDCK cysts, phosphorylation of the ERM protein Ezrin is a major target controlling apical-basal polarity [6]. This regulates where the apical membrane is situated by controlling surface localisation of the apical polarity promoting protein Podocalyxin. In Calu-3, we were unable to detect Podocalyxin expression (unpublished observations), suggesting that alternate Ezrin-binding lumen-promoting proteins may be regulated by ROCK1 phosphorylation. Part of the effect of ROCK1 signalling is to control tensional status of the actin cytoskeleton by controlling phosphorylation and localisation of Myosin-II [7]. Whether Ezrin and Myosin-II are the sole ROCK1 functional targets regulating lumen formation is unclear.

The morphogenetic rearrangements induced by HGF stimulation, and mirrored by ROCK1 inhibition, likely require engagement of the apical polarity programme. Lumen formation in diverse contexts requires a complex interplay between basolateral polarity complexes (Scribble, Discs Large, Lethal Giant Larvae) and the apical complexes (Crumbs/Pals1/PatJ, and Par3-aPKC-Par6-Cdc42) [4]. Mutual antagonism between basolateral and apical complexes ensure that asymmetry between different subdomains of the cell surface can occur [2]. This allows membrane transport to different cortical region [21]. Once established, such cortical asymmetry is self-sustained by positive and negative feedback loops. For instance, in an apical-basal polarized epithelial cell, both apical and basolateral polarity complexes interact with spindle-orientation machinery so that cell divisions occur in the plane of the epithelium, ensuring that cells do not divide into the lumen. The apical component, aPKC, masterminds this process [4]. Interestingly, aPKC and ROCK1/2 exist in antagonism for phosphorylation of Par3 [4]. APKC can phosphorylate Par3, which acts to uncouple Par3 from the Cdc42-aPKC-Par6 complex. However, ROCK-mediated Par3 phosphorylation disrupts aPKC binding to, and phosphorylation of Par3, thereby promoting association of Par3-Par6 [22]. Conversely, aPKC can phosphorylate ROCKs and control their membrane recruitment [23]. The balance between all of these feedback loops controls the orientation of cell division [24]. To induce surface asymmetry, some external cue must tip the balance towards one end of this mutually antagonistic molecular tug-of-war. In Calu-3, this function is performed by HGF/c-Met-dependent ROCK1 inhibition, facilitating basolateral domain formation. ROCK-mediated phosphorylation of Par3 also controls integrin endocytosis [22], and it may control integrin rearrangement to the basolateral domain during multicellular polarisation, similar to control of integrin polarisation in single cells.

In our system, the transition from aggregate to acinus only occurred if cells were cultured in 3D without FCM for at least 4 days. FCM was required for 5 days to induce full polarisation and luminal clearance. Addition of FCM before day 4 resulted in heterogeneous lumen formation, and in the remainder of cases single cell invasion. We hypothesise that polarity rearrangements may only result in a lumen in this system if aggregates are of sufficient size to form a lumen between them, via apoptosis of non-ECM-abutting cells. In the absence of enough cells, invasion may occur. In MDCK cysts, the apical membrane can drive lumen formation or invasion into the ECM [6], depending on the status of the aggregates. A similar biphasic response to ECM cues here appears to occur based on size.

That HGF can induce acinar polarization of Calu-3 lung adenocarcinoma cells is striking, as HGF has been identified as a key factor secreted by lung stroma that promotes the survival of lung cancer cells *in vivo* [25, 26]. Acinar polarization has been shown in other systems to promote resistance to cell death-inducing treatments [27]. What the functional consequence of promoting acinus formation is for treatment for cancer therapies is unknown, but it is tempting to speculate that this may promote survival of lung cancer cells, though this possibility awaits future formal testing.

## 5. Conclusions

In summary, our data indicate conserved ability of a β1-integrin-p190A RhoGAP module in controlling RhoA-ROCK1 signalling for the correct orientation of cell polarity. We now demonstrate that whether this pathway is active in cancer cells may depend on the cellular context and may be controlled in a non-cell autonomous fashion. An unmet need is to understand how to kill residual tumour cells that are resistant to inhibition of cell growth pathways. Our work provides a hint that targeting polarity pathways might be an unrealised, yet effective route to kill residual, resistant cancer cells.

## Acknowledgements

We thank the many investigators that shared reagents. Supported by NIH grants R01DK074398, R01DK091530 and 2P50 GM081879 (KM), and K99CA163535 (DB).

## Author contributions

AD performed the experiments. AD and DB designed experiments and analysed data. DB and ES wrote the manuscript. DB and KM supervised the study.

---

**Supplementary Figure 1: Signalling pathways regulating acinar morphogenesis**.

(A) Aggregates were stimulated with HGF and either PP2 (5 μM), AIIB2 (1:200) or TS2/16 (1:100). Immunoblotting was carried out using anti-pY416-c-Src anti-GAPDH antibodies.

(B) HGF stimulated aggregates were treated with PP2 and stained with anti-pY416-c-Src (green) and Hoechst (blue). Insets show magnified images of pY416-c-Src (middle panels) in untreated (a) or HGF treated (b) samples.

(C) Graph shows the percentage of acini with a single central lumen after treatment with HGF and ROCK inhibition (Y-27632; 10 μM), either alone or in combination.

All graphs are presented as mean percentage +/- s.d. and significance is *p≤0.05, **p≤0.001 and ***p≤0.0001 (n= 3). All scale bars, 50 μm.

## References

1. D.M. Bryant, K. E. Mostov, From cells to organs: building polarized tissue., NatRev Mol Cell Biol 9 (11) (2008) 887–901.

2. E. Rodriguez-Boulan, I. G. Macara, Organization and execution of the epithelial polarity programme, Nat Rev Mol Cell Biol 15 (4) (2014) 225–42.

3. A. W. Overeem, D. M. Bryant, I.S.C. van, Mechanisms of apical-basal axis orientation and epithelial lumen positioning, Trends Cell Biol 25 (8) (2015) 476–85.

4. A. Roman-Fernandez, D. M. Bryant, Complex Polarity: Building Multicellular Tissues Through Apical Membrane Traffic, Traffic 17 (12) (2016) 1244–1261.

5. L. E. O’Brien, M. M. Zegers, K. E. Mostov, Opinion: Building epithelial architecture: insights from three-dimensional culture models, Nat Rev Mol Cell Biol 3(7) (2002) 531–7.

6. D. M. Bryant, J. Roignot, A. Datta, A. W. Overeem, M. Kim, W. Yu, X. Peng, D. J. Eastburn, A. J. Ewald, Z. Werb, K. E. Mostov, A molecular switch for the orientation of epithelial cell polarization, Developmental cell 31 (2) (2014) 171–87.

7. W. Yu, A. M. Shewan, P. Brakeman, D. J. Eastburn, A. Datta, D. M. Bryant, Q. W. Fan, W. A. Weiss, M. M. Zegers, K. E. Mostov, Involvement of RhoA, ROCK I and myosin II in inverted orientation of epithelial polarity., EMBO Rep 9 (9) (2008) 923–9.

8. A. L. Pollack, A. I. Barth, Y. Altschuler, W. J. Nelson, K. E. Mostov, Dynamics of beta-catenin interactions with APC protein regulate epithelial tubulogenesis, J Cell Biol 137 (7) (1997) 1651–62.

9. D. M. Bryant, A. Datta, A. E. Rodríguez-Fraticelli, J. Peränen, F. Martín-Belmonte, K. E. Mostov, A molecular network for de novo generation of the apical surface and lumen., Nat Cell Biol 12 (11) (2010) 1035–45.

10. D. Bryant, K. Mostov, Development: inflationary pressures., Nature 449(7162) (2007) 549–50.

11. Y. Zhu, A. Chidekel, T. H. Shaffer, Cultured human airway epithelial cells (calu-3): a model of human respiratory function, structure, and inflammatory responses, Crit Care Res Pract 2010 (2010).

12. R. Montesano, K. Matsumoto, T. Nakamura, L. Orci, Identification of a fibroblast-derived epithelial morphogen as hepatocyte growth factor, Cell 67 (5) (1991) 901–8.

13. S. K. Mitra, D. D. Schlaepfer, Integrin-regulated FAK-Src signaling in normal and cancer cells, Curr Opin Cell Biol 18 (5) (2006) 516–23.

14. A. E. Bigorgne, H. F. Farin, R. Lemoine, N. Mahlaoui, N. Lambert, M. Gil, A. Schulz, P. Philippet, P. Schlesser, T. G. Abrahamsen, K. Oymar, E. G. Davies, C. L. Ellingsen, E. Leteurtre, B. Moreau-Massart, D. Berrebi, C. Bole-Feysot, P. Nischke, N. Brousse, A. Fischer, H. Clevers, G. de Saint Basile, TTC7A mutations disrupt intestinal epithelial apicobasal polarity, J Clin Invest 124 (1) (2014) 328–37.

15. E. R. Shamir, A. J. Ewald, Three-dimensional organotypic culture: experimental models of mammalian biology and disease, Nat Rev Mol Cell Biol 15 (10) (2014) 647–64.

16. D. K. Marciano, A holey pursuit: lumen formation in the developing kidney, Pediatr Nephrol 32 (1) (2017) 7–20.

17. R. Kalluri, The biology and function of fibroblasts in cancer, Nat Rev Cancer 16 (9) (2016) 582–98.

18. W. Yu, A. Datta, P. Leroy, L. E. O’Brien, G. Mak, T. S. Jou, K. S. Matlin, K. E. Mostov, M. M. Zegers, Beta1-integrin orients epithelial polarity via Rac1 and laminin, Mol Biol Cell 16 (2) (2005) 433–45.

19. L. A. Baker, H. Tiriac, H. Clevers, D. A. Tuveson, Modeling pancreatic cancer with organoids, Trends Cancer 2 (4) (2016) 176–190.

20. L. Julian, M. F. Olson, Rho-associated coiled-coil containing kinases (ROCK): structure, regulation, and functions, Small GTPases 5 (2014) e29846.

21. G. Apodaca, L. I. Gallo, D. M. Bryant, Role of membrane traffic in the generation of epithelial cell asymmetry, Nat Cell Biol 14 (12) (2012) 1235–43.

22. M. Nakayama, T. M. Goto, M. Sugimoto, T. Nishimura, T. Shinagawa, S. Ohno, M. Amano, K. Kaibuchi, Rho-kinase phosphorylates PAR-3 and disrupts PAR complex formation, Developmental cell 14 (2) (2008) 205–15.

23. T. Ishiuchi, M. Takeichi, Willin and Par3 cooperatively regulate epithelial apical constriction through aPKC-mediated ROCK phosphorylation, Nat Cell Biol 13 (7) (2011) 860–6.

24. Y. Hao, Q. Du, X. Chen, Z. Zheng, J. L. Balsbaugh, S. Maitra, J. Shabanowitz, D. F. Hunt, I. G. Macara, Par3 controls epithelial spindle orientation by aPKC-mediated phosphorylation of apical Pins, Curr Biol 20 (20) (2010) 1809–18.

25. K. Matsumoto, M. Umitsu, D. M. De Silva, A. Roy, D. P. Bottaro, Hepatocyte growth factor/MET in cancer progression and biomarker discovery, Cancer Sci 108 (3) (2017) 296–307.

26. R. Straussman, T. Morikawa, K. Shee, M. Barzily-Rokni, Z. R. Qian, J. Du, A. Davis, M. M. Mongare, J. Gould, D. T. Frederick, Z. A. Cooper, P. B. Chapman, D. B. Solit, A. Ribas, R. S. Lo, K. T. Flaherty, S. Ogino, J. A. Wargo, T. R. Golub, Tumour micro-environment elicits innate resistance to RAF inhibitors through HGF secretion, Nature 487 (7408) (2012) 500–4.

27. T. Muranen, L. M. Selfors, D. T. Worster, M. P. Iwanicki, L. Song, F. C. Morales, S. Gao, G. B. Mills, J. S. Brugge, Inhibition of PI3K/mTOR leads to adaptive resistance in matrix-attached cancer cells, Cancer Cell 21 (2) (2012) 227–39.

